# Identification of Factors Complicating Bioluminescence Imaging

**DOI:** 10.1101/511501

**Authors:** Hsien-Wei Yeh, Tianchen Wu, Minghai Chen, Hui-wang Ai

**Affiliations:** Center for Membrane and Cell Physiology, Department of Molecular Physiology and Biological Physics, Department of Chemistry, and the UVA Cancer Center, University of Virginia, 1340 Jefferson Park Avenue, Charlottesville, Virginia 22908, United States of America

## Abstract

*In vivo* bioluminescence imaging (BLI) has become a standard, non-invasive imaging modality for following gene expression or the fate of proteins and cells in living animals. Currently, bioluminescent reporters used in laboratories are mostly derivatives of two major luciferase families: ATP-dependent insect luciferases and ATP-independent marine luciferases. Inconsistent results have been reported for experiments using different bioluminescent reporters and users are often confused when trying to choose an optimal bioluminescent reporter for a given research purpose. Herein, we re-examined inconsistency in several experimental settings and identified factors, such as ATP dependency, serum stability, and molecular size, which significantly affected BLI results. We expect this study will make the research community aware of these factors and facilitate more accurate interpretation of BLI data by considering the nature of each bioluminescent reporter.

## INTRODUCTION

Complementary to magnetic resonance imaging (MRI) and positron-emission tomography (PET), *in vivo* bioluminescence imaging (BLI) offers an accessible and cost-effective method for monitoring gene expression, tumor growth, and protein distribution in animal models.^1-3^ BLI has grown tremendously in the past two decades and become a standard noninvasive and quantitative imaging modality that can provide detailed, time-lapse information on activities and dynamics in animal models without killing animals at various intermediate time points.

Bioluminescent reporters widely used in laboratories can be categorized into two major groups. The first group is mostly derived from insects, such as luciferases from *Photinus pyralis* firefly (FLuc) and click beetles. These enzymes generate light *via* a two-step, oxidative biochemical reaction (Figure 1A). The substrate, D-luciferin, is first adenylated by adenosine triphosphate (ATP) to generate an adenosine monophosphate (AMP)-luciferin intermediate, and subsequently oxidized in the presence of molecular oxygen (O_2_) to yield oxyluciferin in the excited state from which photons with a wavelength of ~ 560 nm are emitted.^4^ The second group comprises ATP-independent luciferases that are cloned from luminous marine organisms.^5^ Coelenterazine (CTZ) is a shared luciferin for a large number of marine luciferases, such as luciferases from *Renilla reniformis* (RLuc),^6^ *Oplophorus gracilirostris* (OLuc),^7^ and *Gaussia princeps* (GLuc).^8^ In the presence of these marine luciferases, CTZ reacts with O_2_ to form a dioxetanone intermediate, which loses CO_2_ to give a high-energy, excited-state coelenteramide (Figure 1B). Subsequently, photons at ~ 480 nm are emitted.^9^

**Figure 1.**
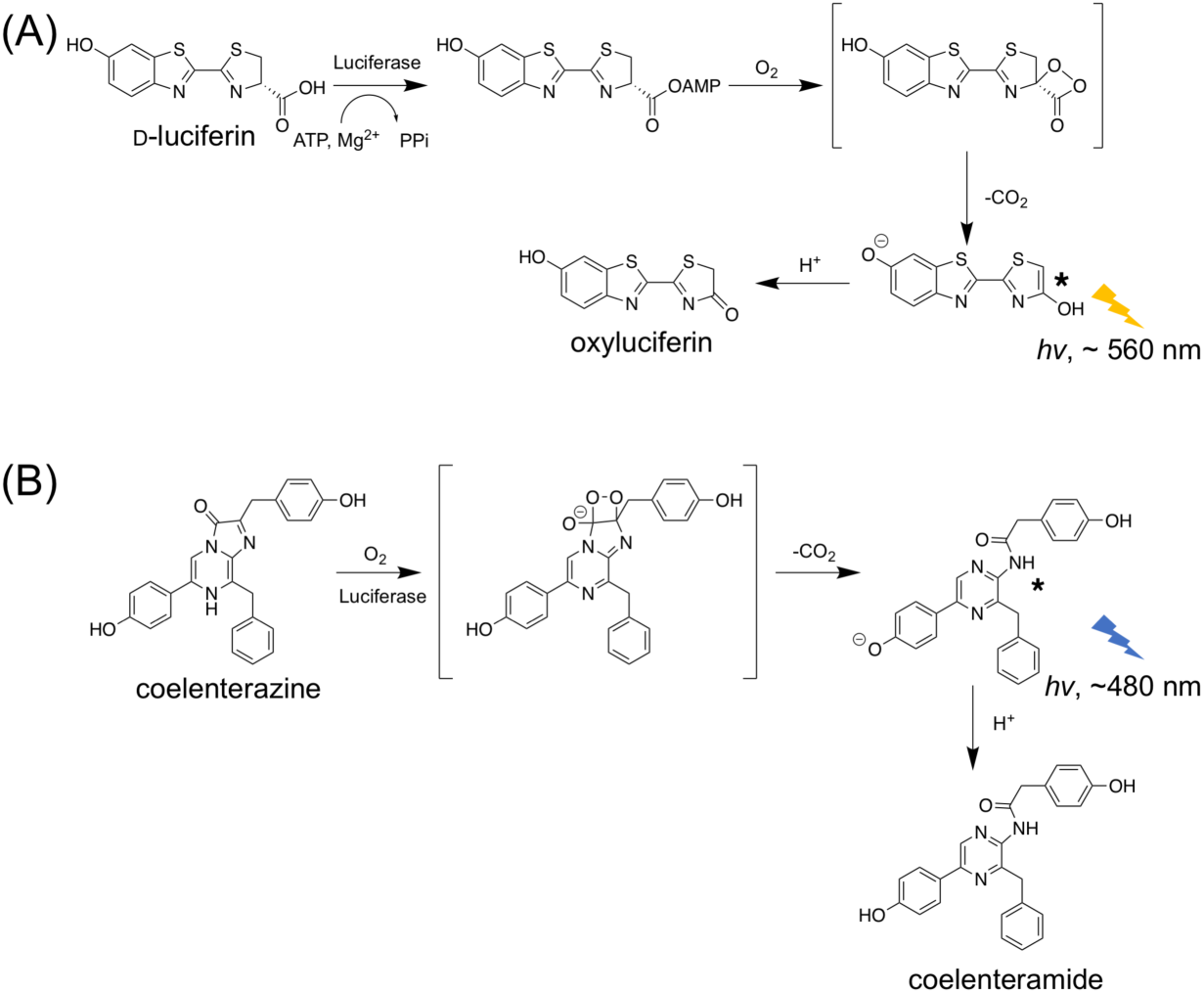
Mechanism of bioluminescence emission generated by luciferase-catalyzed oxidation of (**A**) D-luciferin and (**B**) coelenterazine (CTZ).

FLuc and D-luciferin have been widely utilized for *in vivo* BLI, because its long-wavelength emission is less absorbed and less scattered by mammalian tissue. In recent years, derivatives of FLuc and D-luciferin have been developed and some have generated brighter and even more red-shifted emission. Notably, in early 2018, Iwano *et al*. described an engineered luciferase-luciferin pair, namely Akaluc-AkaLumine, emitting light in the near-infrared (NIR) region for highly sensitive *in vivo* deep-tissue BLI (Figure 2A).^10^

**Figure 2.**
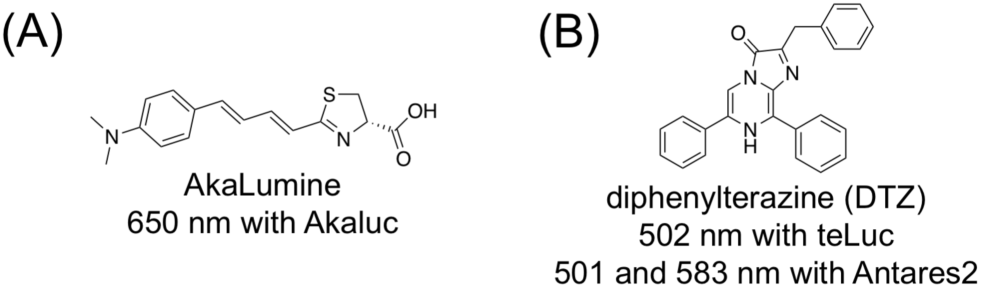
Chemical structures and bioluminescence emission wavelengths of (**A**) AkaLumine and (**B**) diphenylterazine (DTZ).

Conventionally, marine luciferases have limited uses for *in vivo* BLI, because blue photons strongly interact with mammalian tissue, leading to poor light penetration. The recent development of marine luciferase derivatives (*e.g*., NanoLuc and GLuc^11,12^) that produce two orders of magnitude higher photon flux than FLuc has spurred interest in exploring these reporters for *in vivo* BLI. In subcutaneous tumor models,^11,13^ the bioluminescence signal from NanoLuc is comparable to FLuc, despite that FLuc still surpassed NanoLuc for deep-tissue imaging.^14^ Recently, our lab engineered NanoLuc into teLuc, which emits teal photos in the presence of a synthetic diphenylterazine (DTZ) substrate (Figure 2B).^15^ Moreover, further red-shifted emission has been achieved by fusing NanoLuc or teLuc with fluorescent proteins for bioluminescence resonance energy transfer (BRET).^15-18^ These studies have partially fulfilled the increasing demand of using the same reporters for *in vitro* and *in vivo*, fluorescence- and bio-luminescence-based assays. In particular, the Antares2-DTZ pair was able to produce ~10-fold more detectable bioluminescence than FLuc at the standard D-luciferin concentration in a deep-tissue transfection mouse model.^15^

Some researchers have concerns on whether marine luciferase could be used for *in vivo* BLI, because CTZ and its analogs have high internal energies and are prone to oxidation. ^19-21^ In a recent report, the Akaluc-AkaLumine pair was compared with Antares2-DTZ by intravenously (i.v.) injecting Akaluc- or Antares2-expressing HeLa cells into mice via tail vain and further intraperitoneally (i.p.) injecting the corresponding substrates before image acquisition.^10^ The signals from Akaluc-AkaLumine were concentrated to lungs. In contrast, the signals from Antares2-DTZ were stronger but more diffuse than Akaluc-AkaLumine. Subsequently, the diffuse signals from Antares2-DTZ were interpreted as background caused by the auto-oxidation of DTZ. This interpretation is in conflict with our previous observation that direct injection of DTZ into blank mice caused negligible signals, in addition to a plethora of publications which successfully used CTZ or CTZ analogs for *in vivo* BLI. ^17,22-25^

In this manuscript, we re-investigated the performance of Akaluc and Antares2 bioluminescent reporters for tracking i.v.-injected cells in mice, and by using hydrodynamic transfection and xenograft tumor mouse models. We identified several factors, such as ATP dependency, serum stability, and molecular size, which significantly affected BLI results. Our data show that although cautions should be made to maintain the *in vitro* stability of DTZ, background bioluminescence is usually not an issue for its *in vivo* applications. Moreover, our results suggest that it is important to consider the properties of each bioluminescent reporter when interpreting BLI results.

## RESULTS

### DTZ in blank mice generates very low background

We first evaluated bioluminescence background in triplicate by injecting 0.3 μmol of DTZ to BALB/c blank mice that received 2 × 10^4^ untransfected human embryonic kidney (HEK) 293T cells via tail vein. Similar to our previous observation for i.p. injection of DTZ,^15^ the background signal from i.v. injection was also low (Figure 3). Further quantification confirmed that the background signals from i.v. or i.p. injection of DTZ were only slightly above the instrumental background determined from the imaging of untreated blank mice (**Figure S1**, **Supporting Information**). These signals are about two orders of magnitude lower than those of mice injected with luciferase-labeled HEK 293T cells (see below). To confirm that the low background is not strain-specific, we further injected DTZ into B6 albino mice and similar results were observed (Figure 3). Our data collectively suggest that luminescence background from *in vivo* auto-oxidation of DTZ is low and should not be an issue for typical BLI applications.

**Figure 3.**
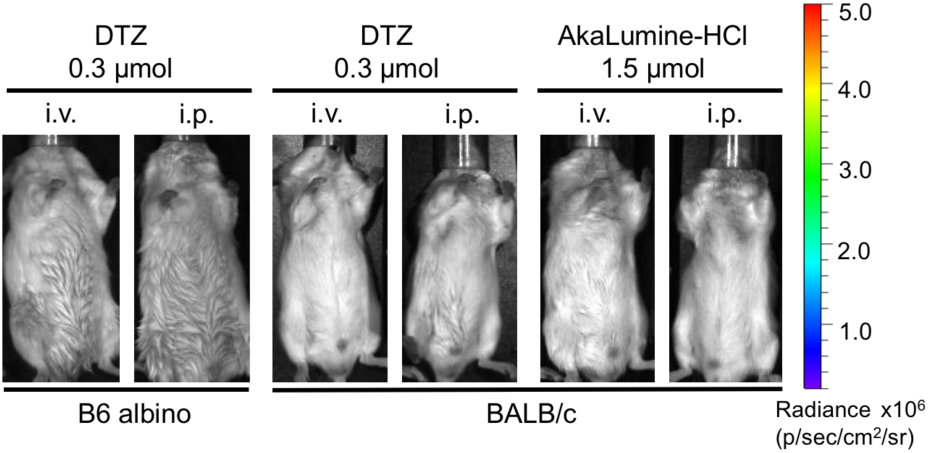
Delivery of DTZ or AkaLumine at the indicated doses via i.v. or i.p. to B6 albino mice or BALB/c mice, yielding negligible background. Luminescence radiance is displayed in the same scale as in Figures 4A and 4B.

### Injection of luciferase-labeled cells into mice releases the luciferase into blood

We next examined the bioluminescence of BALB/c mice at 1, 5, and 24 h after i.v. injection of 2 × 10^4^ HEK 293T transiently expressing Akaluc, teLuc, or Antares2. After i.v. delivery of AkaLumine, we observed bioluminescence signals of Akaluc from the lung area (Figure 4A). This result agrees with previous studies.^10^ In contrast, the signals from teLuc-DTZ were often concentrated in the bladder area; the signals from Antares2-DTZ were observed in the throat, lung, and limbs areas (Figure 4A). Signals from all groups were completely cleared one day after injection of cells to these immunocompetent mice.

**Figure 4.**
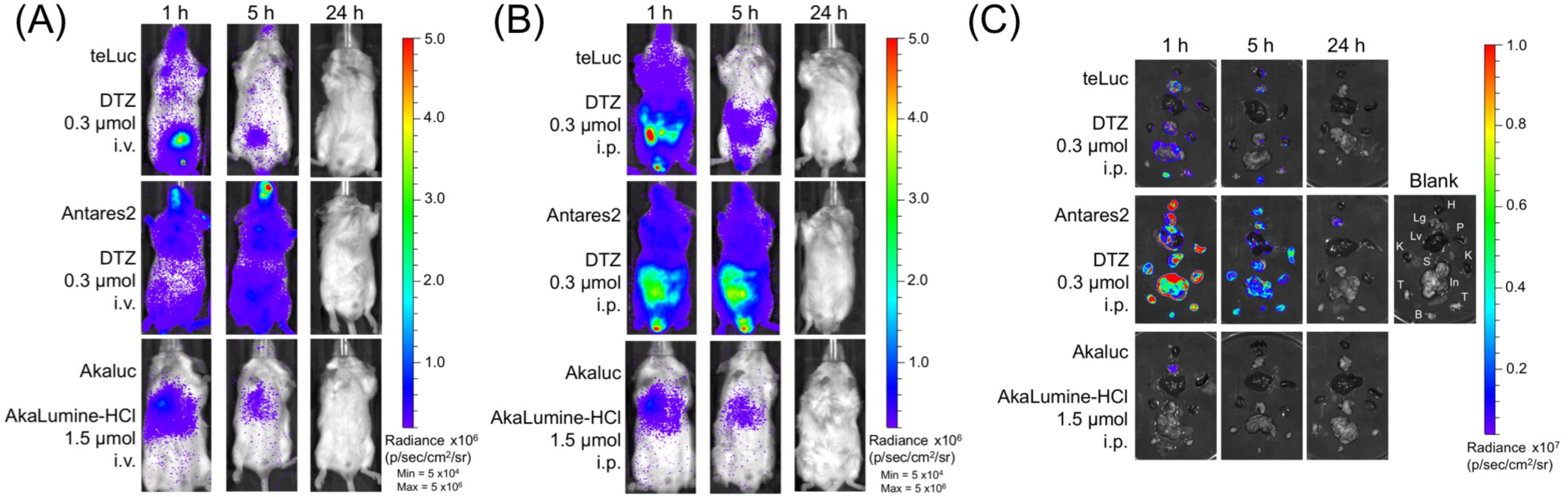
(**AB**) Bioluminescence of mice post i.v. injection of 2 × 10^4^ luciferase-labeled HEK 293T cells (three luciferase-luciferin pairs for comparison: teLuc-DTZ, Antares2-DTZ, and Akaluc-AkaLumine). BALB/c mice (n=3) were imaged 1, 5, and 24 h post injection. Luciferin substrates at indicated doses were delivered via (A) i.v. and (B) i.p., respectively. (**C**) Bioluminescence of extracted organs of mice from panel **B**. H: heart, Lg: lung, Lv: liver, P: spleen, S: stomach, In: intestine, K: kidney, T: testis, and B: bladder.

We further performed BLI after i.p. delivery of corresponding luciferins (Figure 4B). The Akaluc-AkaLumine group had signals in the lung area, similar to the result post i.v. administration of the substrate. Most signals for the teLuc-DTZ group were detected in the abdomen region, while the signals for the Antares2-DTZ group spread through entire mice with peak intensities around the abdomen region.

To further examine signal distribution, mice from each group were sacrificed and their organs were extracted shortly. *Ex vivo* BLI of representative organs showed that that bioluminescence from teLuc-DTZ or Antares2-DTZ were detectable in many organs, whereas bioluminescence for Akaluc-AkaLumine was only observed in lungs (Figure 4C and **Figure S2**). Moreover, we detected strong luminescence from the bladders of mice in the teLuc-DTZ group.

We next collected mouse serum and urine samples for *in vitro* bioluminescence assays. We detected strong teLuc or Antares2 activities from the serums of corresponding mice (Figure 5). Moreover, we observed strong bioluminescence from the urines of mice injected with teLuc-labeled HEK 293T cells. These results suggest that teLuc and Antares2 are released into blood after i.v. injection of their labeled mammalian cells into BALB/c mice. Furthermore, likely due to different molecular sizes, teLuc (19 kDa), but not Antares2 (70.5 kDa), can be excreted into urine via renal filtration.

**Figure 5.**
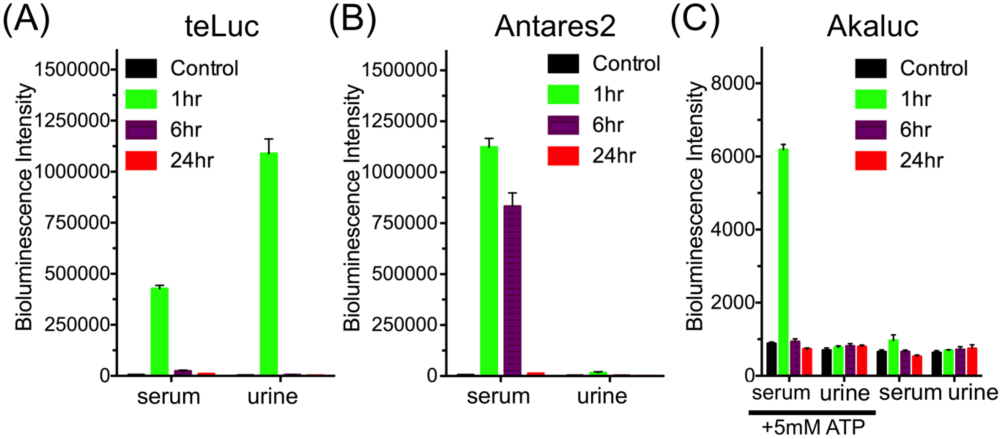
Bioluminescence of serum and urine collected from mice at 1, 6, and 24 h post i.v. delivery of (**A**) teLuc-, (**B**) Antares2-, or (**C**) Akaluc-expressing HEK293T cells, suggesting that luciferases were released into blood and that the small teLuc luciferase further entered urine. Data are presented as mean and s.d. of three biological replicates. In panel C, results are shown for samples with and without supplementing ATP.

We could not detect much bioluminescence directly from the serum and urine samples of mice injected with Akaluc-labeled HEK 293T cells. Since Akaluc is an ATP-dependent luciferase, we next supplemented the samples with 5 mM ATP. In the presence of ATP, bioluminescence was unambiguously detected from the serum, suggesting that Akaluc was also released into blood post i.v. injection of cells.

### In addition to low ATP levels, serum further deactivates Akaluc

The serum bioluminescence of mice in the Akaluc-AkaLumine group was much lower than that of mice in the teLuc-DTZ and Antares2-DTZ groups. This could be explained by the fact that Akaluc is ATP-dependent and that the intrinsic photon flux of Akaluc-AkaLumine is lower than that of teLuc-DTZ or Antares2-DTZ.^10^ To further investigate whether there are other factors, we mixed recombinant teLuc, Antares2, and Akaluc proteins with serum isolated from untreated blank mice and examined luciferase activities in relation to assays in PBS. For the teLuc and Antares2 groups, the total signals in either serum or PBS within the first 10 min post substrate injection were almost identical (Figure 6A), despite that teLuc and Antares2 displayed a flash-type kinetics in PBS but a sustained light output in serum (Figure 6B). In contrast, Akaluc in serum generated very weak bioluminescence, which could be rescued by ~ 4-fold after addition of 5 mM ATP, but still ~ 16-fold lower than its intensity in PBS (Figure 6C and D). These data suggest that, in addition to low ATP levels, mouse serum can further deactivate Akaluc.

**Figure 6.**
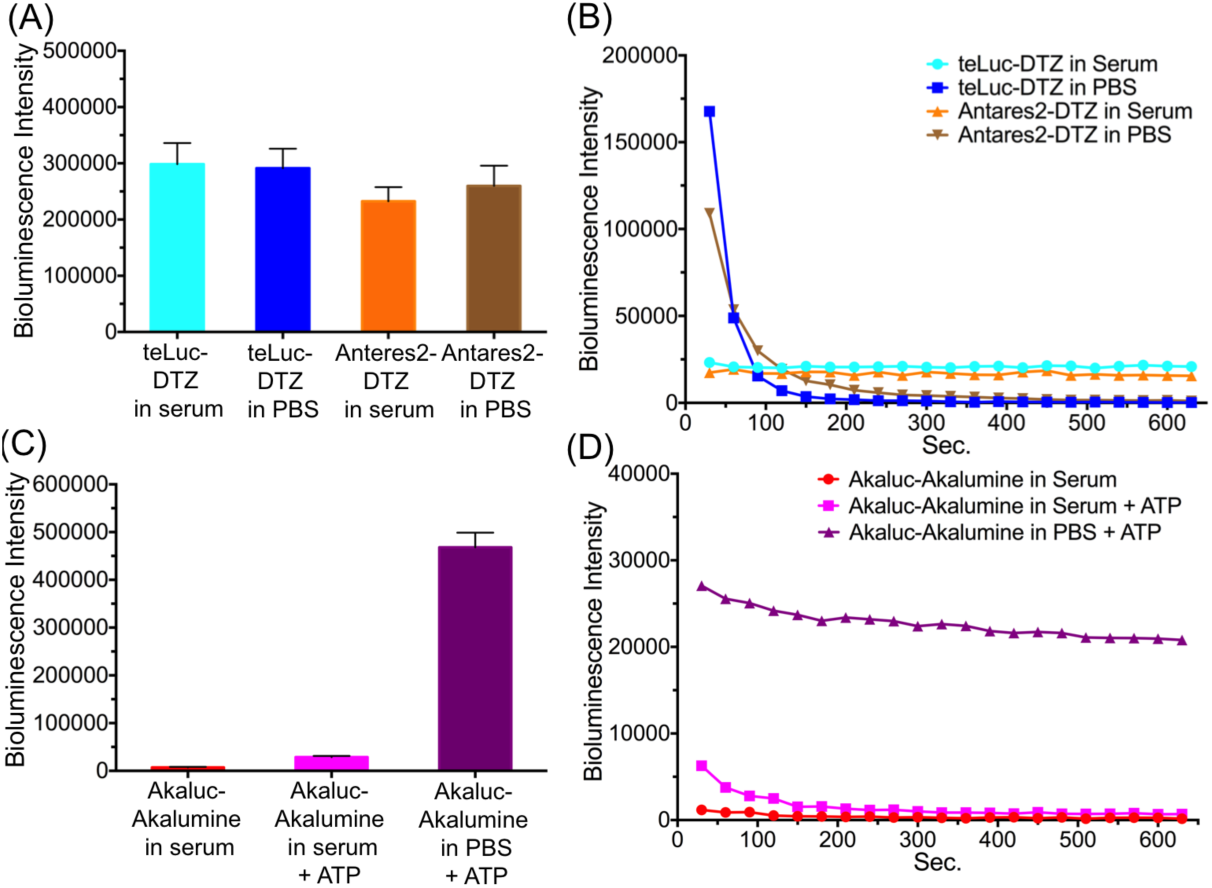
(AB) Bioluminescence intensities (A) and kinetics (B) of purified 1 nM teLuc or Anatres2 in either PBS or mouse serum post injection of 25 μM DTZ. (**CD**) Bioluminescence intensities (C) and kinetic (D) of purified 1 μM Akaluc in PBS or mouse serum post injection of 100 μM AkaLumine and 5 mM ATP.

We thus conclude that luciferases are released into blood after i.v. injection of luciferase-expressing mammalian cells, resulting in diffuse bioluminescence signals from teLuc-DTZ and Antares2-DTZ. We did not observe diffuse bioluminescence signals for Akaluc-AkaLumine, because Akaluc is inactive in blood due to its ATP-dependency and the further deactivation of Akaluc by serum.

### ATP-dependent bioluminescent reporters deplete cellular ATP

Because ATP-dependent luciferases consume ATP to generate bioluminescence, we sought to determine whether these luciferases deplete cellular ATP. We expressed PercevalHR, a previously reported fluorescent ATP/ADP biosensor,^26^in live HEK 293T cells. PercevalHR is an excitation-ratiometric, green fluorescent biosensor with two distinct excitation peaks at ~ 420 and ~ 500 nm. ATP binding increases the excitation of PercevalHR at 500 nm and decreases its excitation at 420 nm.

To our surprise, addition of AkaLumine to mammalian cells generated strong fluorescence (**Figure S3**). AkaLumine itself is only weakly fluorescent (**Figure S4**), but there was obvious, cell-induced fluorescence activation and the resultant fluorescence seemed to accumulate in intracellular vesicles (**Figure S3**). The resultant fluorescent species could be excited with both 405 and 488 nm lasers. Similar phenomena were observed in all tested cell lines, including HeLa, HEK 293T, and SH-SY5Y, and does not require corresponding luciferases since fluorescence was even observed for untransfected cells. Moreover, photoexcitation of AkaLumine-treated cells induced drastic cell morphological changes: the blebbing of cell membrane and the formation of extracellular vesicles were extensive (**Figure S5**). The result suggests that the AkaLumine-induced fluorescent species may act as a photosensitizer.

D-Luciferin is highly fluorescent under 405 nm excitation, but not under 488 nm excitation (**Figures S3** and **S4**). We therefore monitored PercevalHR fluorescence with 488 nm excitation, and by following a previously reported calibration method,^26^ determined ATP level changes in FLuc-expressing HEK 293T cells in response to addition of D-Luciferin. Within a few minutes, the bioluminescence reaction of FLuc-D-luciferin reduced the green fluorescence of PercevalHR, corresponding to a cellular ATP-to-ADP ratio change from > 40:1 to ~ 20:1 (Figures 7). D-luciferin itself did not cause any PercevalHR fluorescence change (**Figure S6A**). Also, the bioluminescence reaction of FLuc-D-luciferin did not affect intracellular pH as monitored by pHRFP, a red fluorescent pH biosensor (**Figure S6B**).^27^

**Figure 7.**
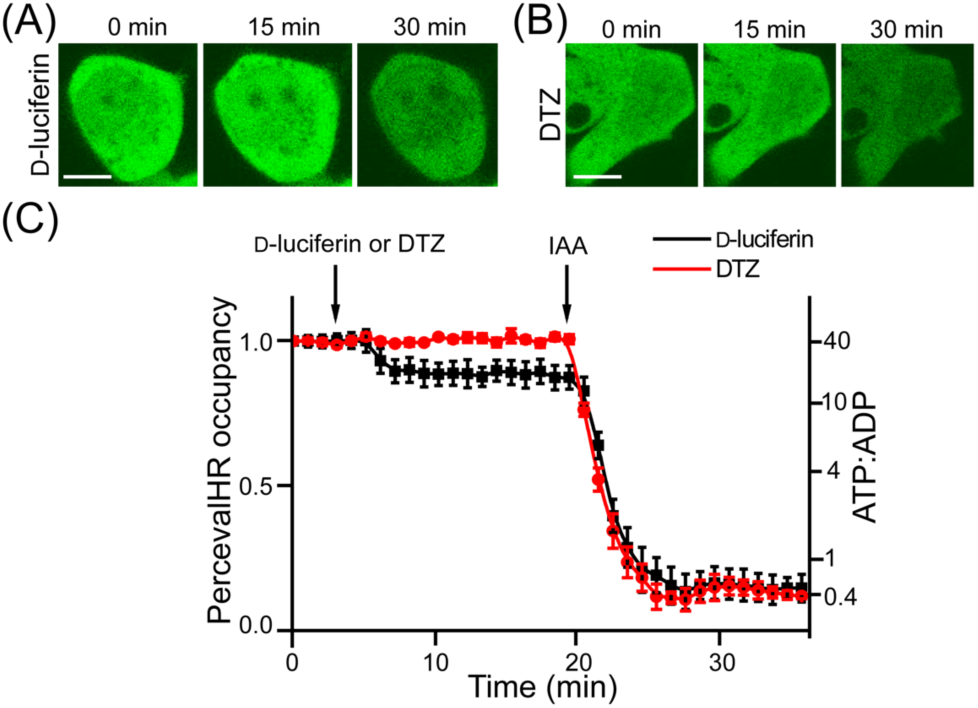
(**AB**) Representative fluorescence images of HEK293T cells expressing PercevalHR and either Fluc (A) or teLuc (B), upon addition of D-luciferin (A) or DTZ (B). Iodoacetic acid (IAA) was used to treat cells after D-luciferin to completely deplete ATP. Scale bars: 10 μm. (**C**) PercevalHR occupancy and estimated ATP:ADP ratio changes upon treatment with D-luciferin or DTZ.

It is worthwhile to note that DTZ does not induce fluorescence in live cells. Moreover, we observed no ATP perturbation from DTZ-treated, teLuc-expressing cells.

### Luciferase is also in the blood of hydrodynamically transfected mice

Hydrodynamic transfection is a systematic gene delivery method.^28^ By rapidly injecting a large volume of DNA to mice via tail vein, this method can lead to transient, high-level gene expression in internal organs, including the liver. Previous studies used FLuc and green fluorescent protein (GFP) to examine the distribution of protein expression after hydrodynamic transfection.^29^We re-investigated this using mice hydrodynamically transfected with Antares2 (Figure 8), which is active in blood. At 10 h after plasmid injection, we observed strong bioluminescence in the abdomen region. We retrieved blood from these mice and detected strong luciferase activity in blood. At 48 h, the bioluminescence became more focused around the liver area, likely due to the activity of Antares2 localized in the liver. The data suggest that after hydrodynamic transfection, proteins encoded by delivered genes are transiently detectable in blood.

**Figure 8.**
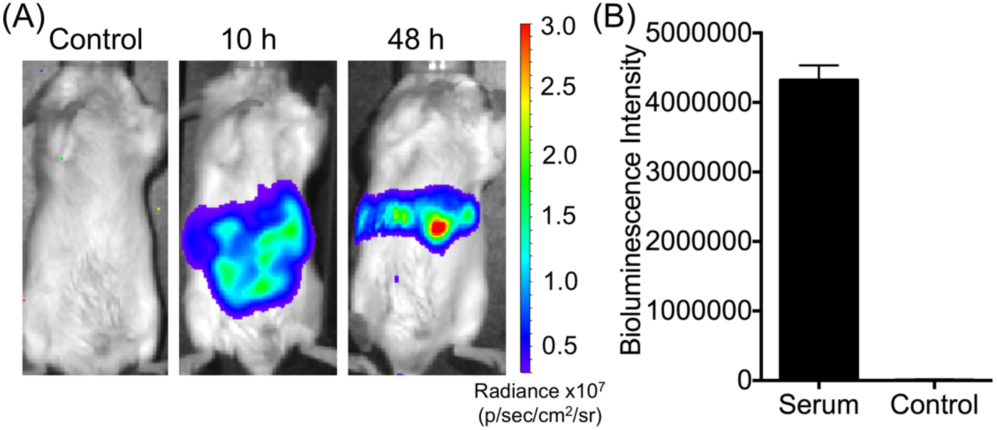
(**A**) BLI of hydrodynamically transfected mice at 10 h and 48 h post transfection. The control mice received hydrodynamic transfection with an empty pcDNA3 plasmid. (**B**) Quantification of bioluminescence of serum isolated from mice at 10 h post hydrodynamic transfection (n=3).

### Luciferases from both families can highlight tumors

BLI has been an important preclinical tool to evaluate the efficacy of a specific treatment by monitor the growth of tumor burden in xenograft models.^3,30^We established HeLa cells that stably express teLuc, Antares2, or Akaluc, and subcutaneously injected 10^4^ or 10^5^cells to the left or right dorsolateral trapezius and thoracolumbar regions of NU/J mice. BLI was performed on day 5 after either i.v. or i.p. administration of corresponding luciferins. On-site implantation of luciferase-expressing tumors gave localized and focused bioluminescence signals no matter ATP-dependent or ATP-independent luciferases were used (Figure 9A). After i.v. administration of the corresponding substrates, the signals for Antares2 were comparable to Akaluc. In addition, we observed ~ 3-fold higher bioluminescence after i.v. administration of DTZ compared to i.p. administration. Moreover, Antares2-DTZ showed better sensitivity than teLuc-DTZ, likely due to its red-shifted emission. Delivery of AkaLumine to Akaluc-expressing tumor xenograft mice via i.v. also yielded higher brightness than i.p. administration, but the difference was within ~ 30% (Figure 9B).

**Figure 9.**
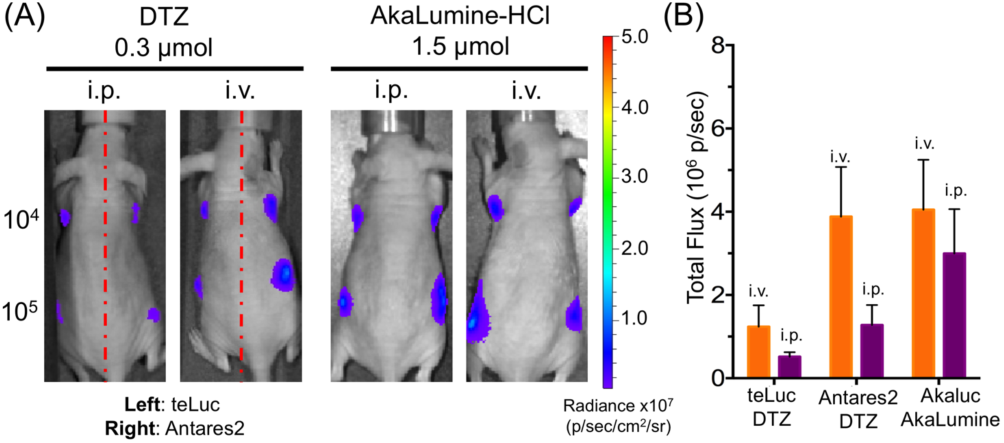
BLI of mice with xenograft tumors. (**A**) BLI (n = 4) on day 5 post injection of 10^4^ or 10^5^ luciferase-expressing HeLa cells to the left and right dorsolateral thoracolumbar regions of NU/J mice. Substrates were administered via either i.v. or i.p. administration. (**B**) Quantitative analysis of bioluminescence intensity of sites inoculated with 10^5^ cells. Data are presented as mean and s.d. of four biological replicates.

## DISCUSSION

In this study, we first investigated reasons for different bioluminescence distributions in mice post injection of luciferase-labeled mammalian cells. Our data support that luciferases were released into blood during or after the injection. Presumably, some of these i.v. injected cells were trapped in lungs,^31^ but some were either lysed during injection or subsequently broken down by the immune systems of these immunocompetent mice. In fact, a few previous studies reported the detection of systemically delivered cells (or in fact their signal labels) in organs other than the lung,^32-37^ corroborating our observations.

The accumulation of teLuc in the bladder suggests that the small size of teLuc makes it permeable to the glomerular basement membrane.^38^ teLuc accumulated in urine still remains enzymatically active. In fact, when mice injected with teLuc-labeled cells were imaged, the bladder area often showed the strongest signals.

For immunocompetent mice injected with Antarea2-labeled cells, signals were observed in the throat, lung, and limb areas. After sacrificing these mice, bioluminescence was detectable in blood and many internal organs. Since bioluminescence is attenuated approximately by a factor of 10 for every centimeter depth of tissue^39^, we suspect that those signals from superficial blood or the lymphatic system around the throat may partially overwrite signals from deep tissues.

ATP-dependent luciferases such as Akaluc is inactive in blood because of relatively low ATP levels in the extracellular space and deactivation of Akaluc by blood. Only signals from the lung were observed for mice injected with Akaluc-labeled cells, because the lung trapped live mammalian cells, which can produce ATP to support the bioluminescence reaction of Akaluc. In contrast, dead cells or Akaluc released in the circulation systems were not detectable when whole mice were imaged.

Although ATP-dependent luciferases, such as FLuc, have been routinely used, their impact on cellular ATP levels has not yet carefully characterized. Assuming the bioluminescence quantum yield is 0.4,^40^ to produce each photon, 2.5 ATP molecules are consumed. This is a significant metabolic burden, considering that the luciferases may reach micromolar concentrations in live cells. We co-expressed PercevalHR, a fluorescent biosensor for monitoring the ATP-to-ADP ratio, with FLuc and observed the decrease of the intracellular ATP-to-ADP ratio from > 40:1 to ~ 20:1 during the bioluminescence reaction between FLuc and D-luciferin. We want to note that our analytical method is limited since PercevalHR is saturated when ATP-to-ADP ratios are more than 40:1. Some previous studies in fact suggest that, the ATP-to-ADP ratio could be up to ~ 200:1 in healthy cells.^41^ This metabolic disruption issue of ATP-dependent luciferases casts doubt on whether they perturb biology. Moreover, because the ATP level is a variable in biological systems and the availability of ATP may sometimes become a limiting factor, ATP-dependent bioluminescence systems may actually report the co-exist of ATP, the substrate, and the luciferase.

ATP-independent luciferases, such as teLuc and Antares2, can overcome the ATP-dependency issue of ATP-dependent luciferases. However, current ATP-independent bioluminescence systems have a different set of caveats.

First, despite that the luminescence background caused by DTZ auto-oxidation is low and not a problem for most applications, the background might become an issue for detection of very low signals, such as bioluminescence from single cells. We suggest that researchers should always include suitable controls for their particular experiments to exclude autoluminescence. Also, we envision that future research may lead to new CTZ and DTZ analogs to greatly enhance their stability and reduce their auto-oxidation, thereby broadening the application boundary of marine luciferases.

Second, DTZ and its analogs have a delayed biodistribution, as demonstrated by us (Figures 4A, **4B**, and **9B**) and others.^42,43^ The route of administration may have to be optimized for each type of experiments.^23,44^ In our xenograft tumor model, i.v. administration was clearly more effective than i.p. delivery of the substrate. We want to note that AkaLumine may also have uneven biodistribution, since it seemed to be accumulated in the liver as shown by our near-infrared (NIR) fluorescence imaging of organs of AkaLumine-injected mice (**Figure S7**).

Moreover, DTZ and its analogs have relatively poor water solubility, resulting in limited *in vivo* delivery dosages. Although these ATP-independent luciferases can generate much higher photo flux than FLuc and Akaluc *in vitro*, their brightness *in vivo* is compromised by the low accessibility of substrates and their relatively short-wavelength emission (see below). We expect that further chemical modifications of DTZ and its analogs will improve their water-solubility, biocompatibility, and biodistribution.

Furthermore, the emission of marine luciferase derivatives is less red-shifted than the emission of FLuc or Akaluc, leading to lower tissue penetration. In particular, the depth and opacity of the tissues will complicate signal acquisition and data interpretation. Signals from superficial targets may dominate signals from deep tissue. Also, because BLI is intrinsically a 2-dimemsional imaging modality, light transmission is dependent of mouse strains. For example, the interaction of light with black mice, albino mice, or hairless mice would be very different.^45^ Because of these reasons, we recommend the capturing of bioluminescence on both ventral and dorsal sides. Moreover, to increase tissue penetration, we expect that further studies will continuously red-shift the emission of marine luciferase derivatives.

In this study, we also investigated hydrodynamic transfection of mice with Antares2. We detected high bioluminescence in blood. It has been reported that high intravascular pressure can stretch the structure of hepatocytes, and subsequently lead to whole-body re-distribution of injected plasmids.^46^ This disruption seemed to be effective in releasing of newly synthesized Antares2 to blood. Previous studies used FLuc and GFP as the reporters to study gene expression across various organs, but FLuc and GFP were unable to provide information on gene expression in blood.^29^ Therefore, ATP-independent Antares2 provides an alternative insight to this process.

We further demonstrated that both ATP-dependent Akaluc and ATP-independent Antares2 could be used to detect xenograft tumors. Bioluminescence signals were observed at sites for tumor implantation. This result strongly supports that ATP-independent Antares2 does not have intrinsic tendency to generate diffuse signals. Antares2 caused diffuse signals in some systematic delivery experiments, only because the luciferase indeed entered and were still active in body fluids.

Because of the ATP-independency of Antares2, we expect it may complement ATP-dependent luciferases, such as FLuc and Akaluc, and may reveal new biological insights not detectable with ATP-dependent luciferases. In addition, we deem the use of dual BLI with both ATP-dependent and ATP-independent luciferases, not only for tracking multiple targets, but also for factual reporting of a single biological process by breaking natural limitations (*e.g*., substrate biodistribution, light attenuation by tissue, and ATP availability) of a single bioluminescence system.

Although bioluminescent reporters have been discovered and further developed in laboratories for quite long time, they are far from ideal for biological applications. This paper has identified several key issues for common ATP-dependent and ATP-independent reporters. These issues complicate BLI results. Therefore, in order to accurately interpret the data, one has to consider the chemical, biological, and photophysical nature of each bioluminescent reporter. Further effort is urgently needed to make these reporters more robust and biocompatible and to simplify BLI experiments and result interpretation.

## EXPERIMENTAL METHODS

### Mammalian cell culture and transfection

HEK 293T cells were cultured at 37 °C with 5% CO_2_ in Dulbecco’s Modified Eagle’s Medium (DMEM) supplemented with 10% fetal bovine serum (FBS). Transfection mixtures were prepared with 3 μg of plasmid DNA and 9 μg of PEI (polyethylenimine, linear, MW 25 kDa) in DMEM and incubated for 20 min at room temperature. After removing cell culture media, transfection mixtures were added to cells at ~ 70% confluency on 35-mm culture dishes seeded 1 d before transfection. Incubation lasted for 3 h at 37 °C. Fresh DMEM containing 10% FBS was used to replace the transfection mixtures. Cells were cultured at 37 °C in a CO_2_ incubator for another 24 h before use.

### Bioluminescence imaging of i.v. injected HEK 293T cells

BALB/c, NU/J and B6 albino mice were obtained from the Jackson Laboratory (Cat. No. 000651, 002019 and 000058). Animals were maintained and treated in standard conditions that complied with all relevant ethical regulations. All animal procedures were approved by the UVA Institutional Animal Care and Use Committee. HEK 293T cells expressing teLuc, Antares2, or Akaluc were trypsinized, pelleted, and re-suspended in 100 μL PBS. Cell numbers were determined using a hemocytometer. 20000 cells were injected into mice placed in a restrainer via tail vein. Mice were allowed to recover. Bioluminescence images were taken at 1h, 5h, and 24h post injection of cells. Mice anaesthetized using isoflurane were i.p. or i.v. injected with 0.3 μmol DTZ in a 100 μL solution containing 8% glycerol, 10% ethanol, 10% hydroxypropyl-β-cyclodextrin, and 35% PEG 400 in water or 1.5 μmol AkaLumine-HCl in 100 μL saline. The 0.3 μmol DTZ solution was each time made fresh from a DTZ (50 mM) stock solution in 1:1 (v/v) EtOH and propylene glycol containing 10 mM HCl (stored at −80°C). Mice under isoflurane anesthesia on a heat pad were next immediately imaged with a Caliper IVIS Spectrum over a course of 20 min. The following conditions were used for image acquisition: open filter for total bioluminescence, exposure time = 10 s, binning = small, field of view = 21.6 × 21.6 cm, and f/stop = 1. Image analysis was performed using the Living Image 4.3 software. After imaging at each time point, one mouse in each group was sacrificed. Organs were harvest and plated on a petri dish and bioluminescence images were acquired immediately with 1-s exposure. Blood and urine samples were collected for further evaluation. **Bioluminescence assays of blood and urine.** After collection, blood was left to clot at room temperature for 30 min. Clots were removed by centrifugation at 2000xg for 10 min. The resultant supernatant (a.k.a serum) was store at −80°C. 5 μL of urine or serum was diluted in 100 μL PBS to a white 96-well plates (Costar 3912). 100 μL of DTZ was added to a final concentration of 25 μM. Bioluminescence activities were immediately measured on a Synergy Mx Microplate Reader (BioTek). To monitor Akaluc activity in serum or urine, the final concentration of AkaLumine-HCl was at 100 μM. Moreover, 5 mM ATP and 5 mM MgSO_4_ were supplemented as needed.

### Fluorescence imaging of ATP in HEK 293T cells

HEK 293T cells were cultured and transfected as aforementioned. Images were acquired on a Leica DMi8 inverted microscope equipped with the SPE confocal module. Cells were cultured in DMEM (no phenol red) with 4.5 g/mL glucose. A 488 nm laser was used to excite PercevalHR and emission was collected from 510 nm to 600 nm. Intervals between each image were 5 second. 1 mM D-luciferin or 50 μM DTZ was added to cells. For internal calibration, iodoacetic acid (IAA) was added to a final concentration of 5 mM after D-luciferin to completely deplete ATP.^26^

### Hydrodynamic transfection of BALB/c mice

20 μg of each luciferase-expressing plasmid in 2 mL sterilized saline was injected into restrained mice via tail vein over 8 s. Mice were allowed to recover and monitored until their breathing rate returned to normal. BLI was performed 10 and 48 h post transfection.

### NU/J xenograft model

HeLa cells stably expressing luciferases were dissociated with trypsin and re-suspended in 10 mL DMEM. Cell numbers were determined using a hemocytometer, and cell viability was determined using a trypan blue exclusion test. 10^4^ or 10^5^ cells were re-suspended in 100 μL FBS-free DMEM containing 50% Matrigel matrix (Corning). 8-week-old female nude mice were first anesthetized using isoflurane. Cells were subcutaneously injected into the left and right dorsolateral trapezius regions or thoracolumbar regions. Mice were recovered on heat pads for 5 min while cells were allowed to settle. BLI were performed on day 5 post cancer cell inoculation.

## AUTHOR INFORMATION

### Notes

H.w.A. and H.W.Y. are coinventors of a patent filed by the Regents of the University of California, titled “Red-Shifted Luciferase-Luciferin Pairs for Enhanced Bioluminescence”. The sequences of teLuc, Antares2, and Akaluc can be accessed from NCBI GenBank under accession numbers KX963378, KY474379, and LC320664.

## ABBREVIATIONS

BLI: bioluminescence imaging
MRI: magnetic resonance imaging
PET: positron-emission tomography
FLuc: firefly luciferase
BRET: bioluminescence resonance energy transfer
GFP: green fluorescent protein

