## Supplementary information for "Identification of Factors Complicating Bioluminescence Imaging"

### Comparison of Luciferases for Bioluminescence Imaging

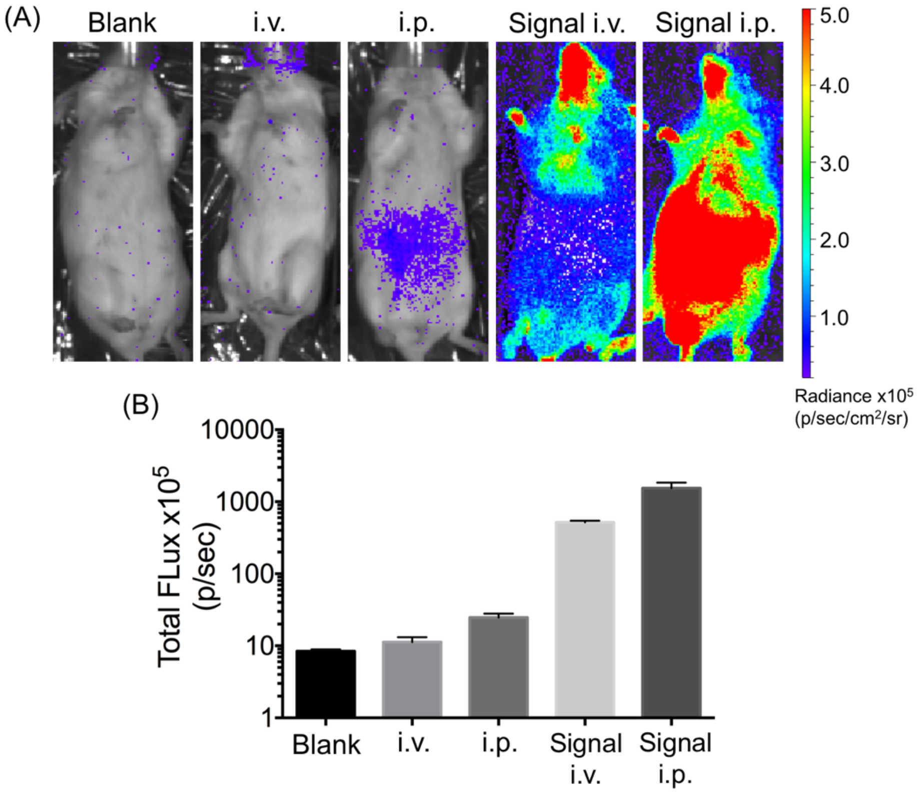


**Figure S1**. (**A**) Representative bioluminescence images of BALB/c mice (from left to right: blank, i.v. injection of DTZ, i.p. injection of DTZ, and i.v. or i.p. injection of DTZ after i.v. injection of Antares2-expressing HEK 293T cells). To show the existence of background signals, luminescence radiance is displayed in a compressed scale. (**B**) Quantitative analysis of signals integrated over areas. The y axis is in a logarithmic scale. Data are presented as mean and s.d. of three biological replicates.

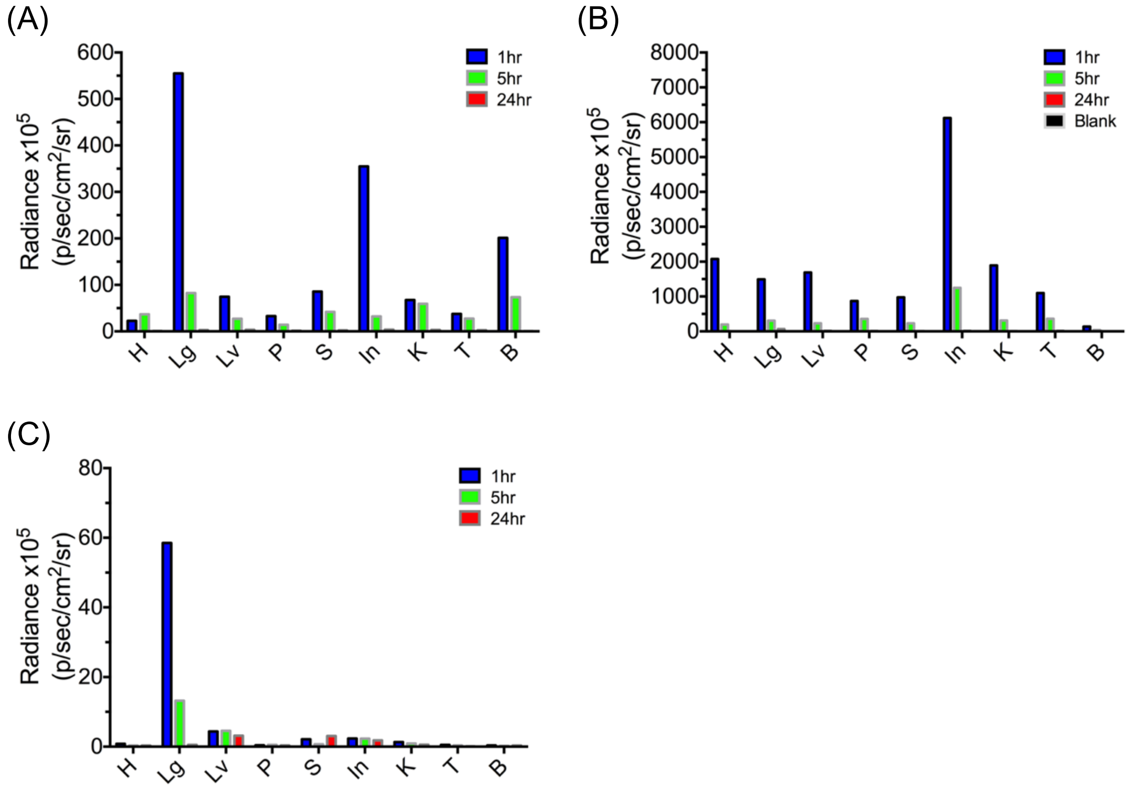


**Figure S2**. *Ex vivo* bioluminescence of organs harvested from mice at 1, 5, 24 hours after i.v. injection of HEK293T cells expressing (**A**) teLuc, (**B**) Antares2, or (**C**) Akaluc (n = 1 for each group). H: heart, Lg: lung, Lv: liver, P: spleen, S: stomach, In: intestine, K: kidney, T: testis, and B: bladder.

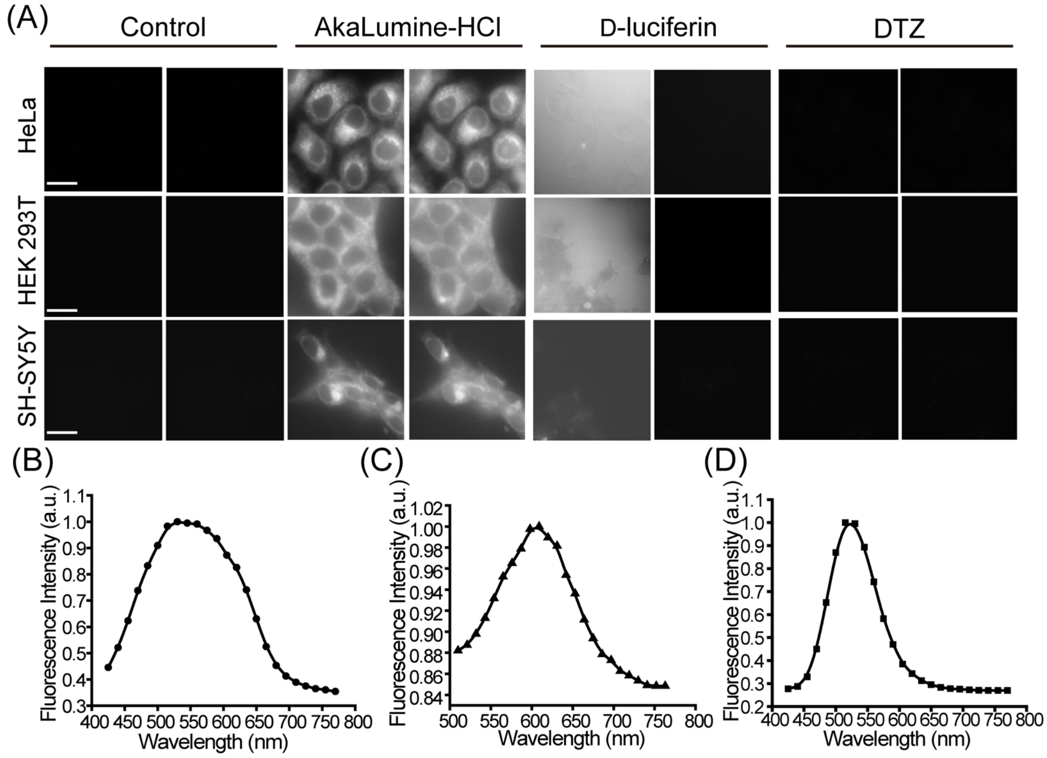


**Figure S3.** (**A**) Fluorescence imaging of live mammalian cells after incubation with indicated luciferins for 10 min. 100 μM AkaLumine-HCl, 200 μM D-luciferin, or 50 μM DTZ was added to cell lines, including HeLa, HEK 293T, and SH-SY5Y. These concentrations were chosen since each compound has different cell permeability and previous publications used these concentrations for corresponding assays.[1-4] Images were captured with 405-nm or 488-nm laser excitation, and emission was collected from 510 nm to 600 nm (scale bars: 20 μm). (**BC**) Emission spectra of AkaLumine-treated HEK 293T cells with 405 nm (B) or 488 nm (C) laser excitation. (**D**) Emission spectrum of D-luciferin-treated HEK 293T cell with 405-nm laser excitation.

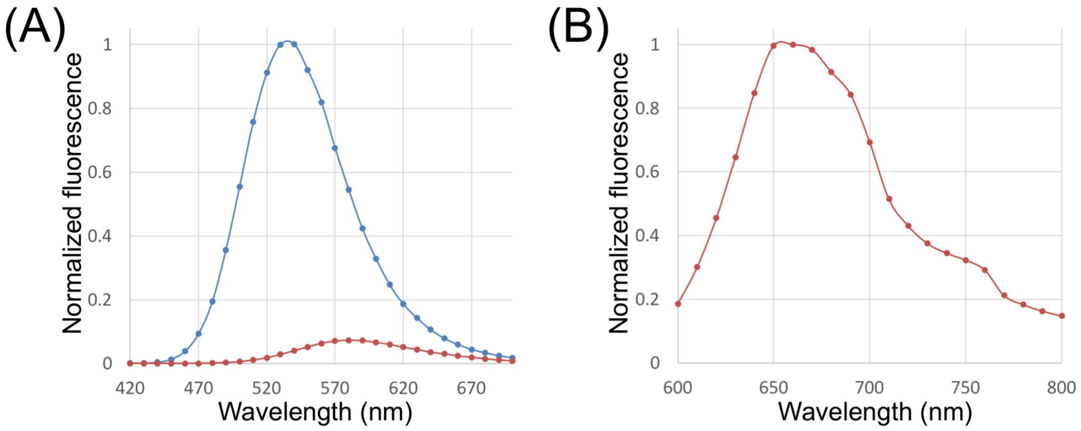


**Figure S4.** (**A**) Fluorescence emission of 1 mM D-luciferin (blue) or AkaLumine (red) in PBS with 400 nm excitation. (**B**) Fluorescence emission of 1 mM AkaLumine (red) in PBS with 580 nm excitation.

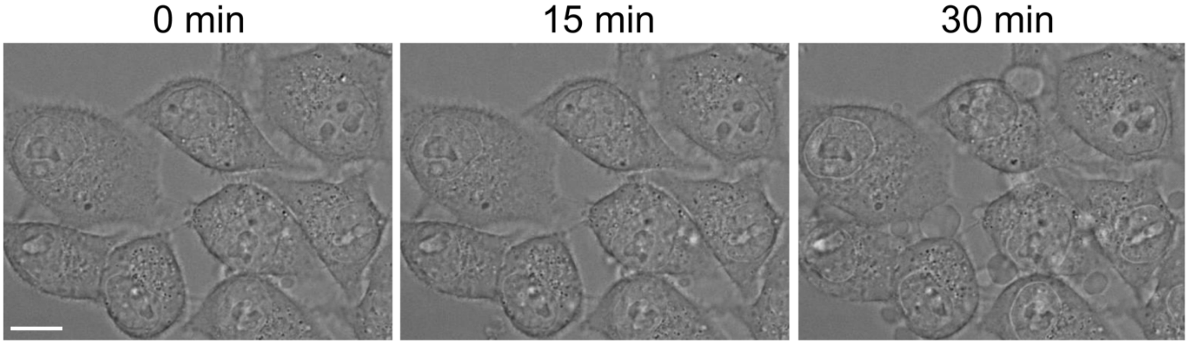


**Figure S5.** Morphological changes of HEK 293T cells treated with 100 μM AkaLumine and imaged with 488-nm laser scanning, showing the blebbing of cell membrane and the formation of extracellular vesicles.

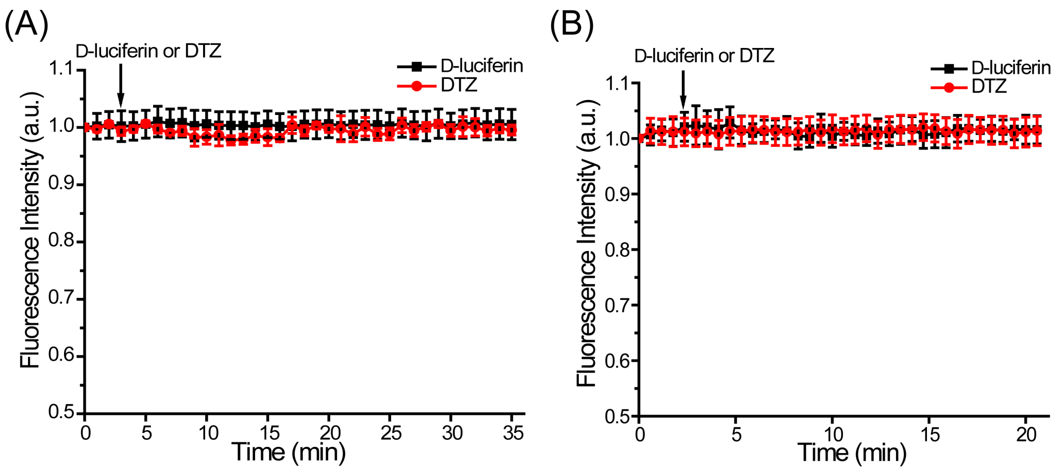


**Figure S6.** (**A**) PercevalHR fluorescence in HEK 293T upon treatment with D-luciferin or DTZ in the absence of FLuc or teLuc, showing no change in ATP occupancy. (**B**) pHRFP fluorescence in HEK 293T upon treatment with D-luciferin or DTZ in the presence of the corresponding luciferase, showing that the corresponding bioluminescence reactions do not affect intracellular pH.

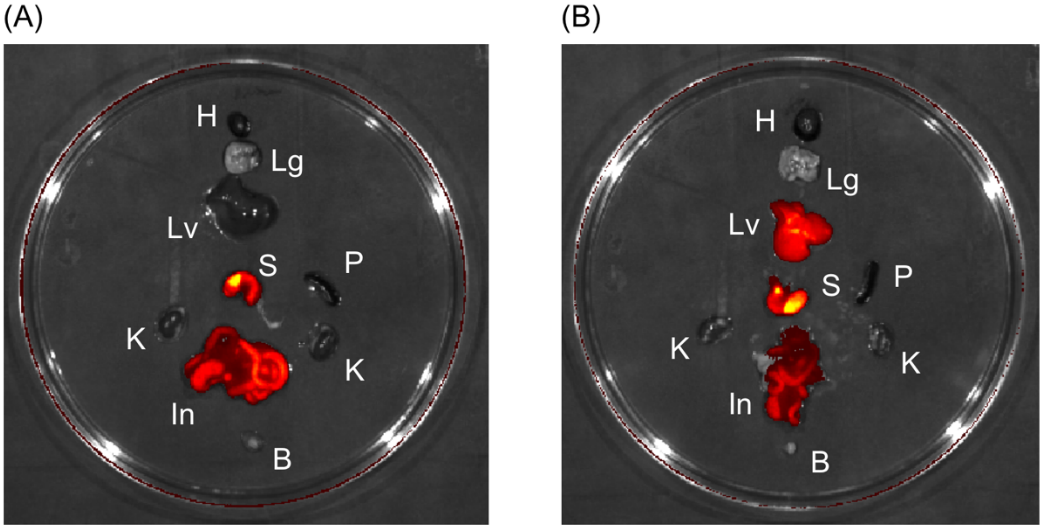


**Figure S7.** Fluorescence imaging (excitation: 605/25 nm bandpass; emission: 660/25 nm bandpass) of organs harvested from (**A**) an untreated BALB/c mouse, (**B**) a BALB/c mouse i.v. injected with 1.5 μmol AkaLumine-HCl. Near-infrared (NIR) background fluorescence, mostly in the gastrointestinal tract, was observed for mice, because chlorophyll from plant-based diet (e.g., alfalfa) is fluorescence in this spectral range.[5] In addition to the background, the AkaLumine-injected mouse showed high NIR fluorescence in the liver. H: heart, Lg: lung, Lv: liver, P: spleen, S: stomach, In: intestine, K: kidney, and B: bladder.
